# Distributional context reweights duration-related activity across timing-related regions

**DOI:** 10.64898/2026.05.12.724553

**Authors:** Cemre Baykan, Si Cheng, Zhuanghua Shi

**Author notes:** corresponding author: Cemre Baykan, Philipps-Universität Marburg, Fachbereich Psychologie, AG Sensomotorisches Lernen, Marburg, Hessen, Germany, or via. This project was supported by DFG project SH 166/10-1 to Z.S. and DAAD scholarship 57440921 to C.B.

## Abstract

Duration judgments are calibrated by the statistics of recent temporal experience, but how this contextual influence is expressed across human timing-related brain regions remains unclear. We used functional MRI while 24 participants performed a visual temporal bisection task under two distributional contexts: one biased toward longer durations and one biased toward shorter durations. Behaviorally, distributional context shifted subjective duration judgments without changing in temporal precision, consistent with a prior-related bias. Neurally, duration-related BOLD responses varied across a priori timing regions as a function of context. The clearest region-specific evidence was stronger left-insula duration modulation in the short-biased context. SMA/pre-SMA contributed to the broader timing-network pattern, and its context-specific asymmetry was sensitive to the alignment of the duration regressor. Exploratory SMA-seeded connectivity analyses provided converging evidence that context also modulated duration-related coupling with timing- and attention-related regions. Together, these findings indicate that distributional context impacts the relative weighting of established timing-related regions during duration judgments, rather than recruiting anatomically separate timing systems.

## Introduction

Rather than encoding every element of a perceptual stream independently, the brain extracts summary statistics that distill recent input into a compact, actionable representation (Ariely, 2001; Chong & Treisman, 2003; Haberman & Whitney, 2007). This ensemble-coding principle extends to time perception. Duration judgments are assimilated to the statistical average of recent experience, and that assimilation produces central-tendency and contextual biases (Glasauer & Shi, 2022; Jazayeri & Shadlen, 2010; Shi et al., 2013). In temporal bisection, observers classify intervals relative to learned short and long anchors, yet their judgments also track the distribution of recently encountered durations: long-biased contexts shift the point of subjective equality (PSE) toward longer durations, and short-biased contexts shift it toward shorter durations (Baykan et al., 2023; Zhu et al., 2021). Bayesian and predictive-coding accounts explain such biases as the integration of sensory evidence with a distributional prior (Petzschner et al., 2015; Shi et al., 2013). The ensemble-distribution account goes further: the prior guiding categorical timing decisions reflects not only the mean of recent durations but also their variance and skew (Zhu et al., 2021). The behavioral phenomenon itself is well established; what remains open is how a distributional prior is expressed within the human timing system.

Duration processing engages a distributed cortico-subcortical network rather than a single timing module (Paton & Buonomano, 2018). Meta-analytic and task-based neuroimaging studies implicate SMA/pre-SMA, the putamen, the cerebellum, the inferior frontal gyrus (IFG), the insula, and parietal cortex across sub-second and supra-second timing tasks (Coull et al., 2004; Mondok & Wiener, 2022; Nani et al., 2019; Wiener et al., 2010). SMA/pre-SMA activity has been linked to elapsed time, temporal working memory, motor simulation, and anticipation (Coull et al., 2016; Merchant et al., 2013). The putamen is central to cortico-striatal timing theories, including the beat-frequency model, in which the striatum reads out duration from coincident cortical oscillations (Matell & Meck, 2004). The cerebellum is consistently implicated in sub-second timing, whereas IFG and parietal regions contribute to temporal attention and decision demands (for reviews, see Mondok and Wiener 2022; Nani et al. 2019). The insula, in turn, has been tied to subjective duration, salience, and interoceptive timing (Craig, 2009; Wittmann, van Wassenhove, et al., 2010). These regions are unlikely to be recruited in all-or-none fashion by temporal context. A more plausible account is that distributional context alters the relative weighting, gain, or detectability of duration-related signals within this established network.

Several lines of evidence suggest that temporal context can modulate this network. Electrophysiological studies show that distributional context affects both anticipatory and post-stimulus timing signals. The contingent negative variation (CNV) and the late positive component of timing (LPCt) each shift with context, implicating both accumulation and decision stages (Baykan et al., 2023; Baykan & Shi, 2023; Damsma et al., 2021; Ng et al., 2011; Ofir & Landau, 2022; van Rijn et al., 2011). Causal evidence points to medial frontal and premotor regions: stimulating SMA changes the precision or the bias of temporal bisection (Méndez et al., 2017; Wiener et al., 2018), and dorsal premotor cortex contributes to temporal prediction (Lazzari et al., 2025). A further set of findings ties SMA/pre-SMA to the prior more directly: SMA activity can track the cumulative hazard carried by the prior rather than elapsed time alone (Cui et al., 2009), primate pre-SMA population dynamics are warped by the sampled distribution (Sohn et al., 2019), and pre-SMA neurons encode adaptive category boundaries that move with context (Mendoza et al., 2018).

Behavioral and EEG studies thus establish that distributional context changes temporal decisions and the slow neural dynamics that precede and follow them, but they cannot localize the effect within timing-related circuitry. Functional MRI offers that spatial leverage. It can test whether contextual modulation takes the form of differential weighting across established timing regions (SMA/pre-SMA, putamen, IFG, cerebellum, and insula). The question matters beyond anatomical localization. If identical target durations are processed differently depending on the sampled distribution, a timing model must explain not only how duration is encoded but also how a distributional prior reweights the contributions of these regions to the final judgment. Most prior fMRI timing studies have examined a single context, leaving them poorly suited to detect the context-by-duration interactions through which ensemble statistics modulate duration-related neural signals.

Here we tested whether distributional context modulates duration-related BOLD activity within established timing regions during visual temporal bisection. Twenty-four participants judged identical target durations under a long-biased and a short-biased frequency distribution. Guided by the prior work (Zhu et al., 2021), we asked two questions. First, do duration-related BOLD responses vary with context across a priori timing ROIs, treating SMA/pre-SMA as a literature-motivated candidate region? Second, does SMA-centered functional coupling during duration processing vary with context? These regional and connectivity analyses together address how and where a distributional prior is expressed within the human timing network.

## Method

### Participants

Twenty-four right-handed adults (13 female; age range 21–36, mean 26.2 years) with normal or corrected-to-normal vision and no history of neurological or psychiatric disorders took part. Sample size was determined a priori from the ensemble context effect in Zhu et al. (2021; η_g_ = 0.26); a power analysis (α = .05, 1 − β = .95) required a minimum of 16 participants, and we recruited 24 to buffer against motion-related loss. All participants gave written informed consent, received 15 EUR/h, and were naive to the study purpose. The study was approved by the Ethics Board of the Department of Psychology, LMU Munich (29.05.2018).

### Stimuli and Procedure

Participants performed a visual temporal bisection task (Figure 1) in which they categorized target durations as “short” or “long” relative to previously learned anchors, with temporal context manipulated by the frequency distribution of target durations across two conditions: ascending frequency (AF, long-biased) and descending frequency (DF, short-biased).

**Figure 1.**
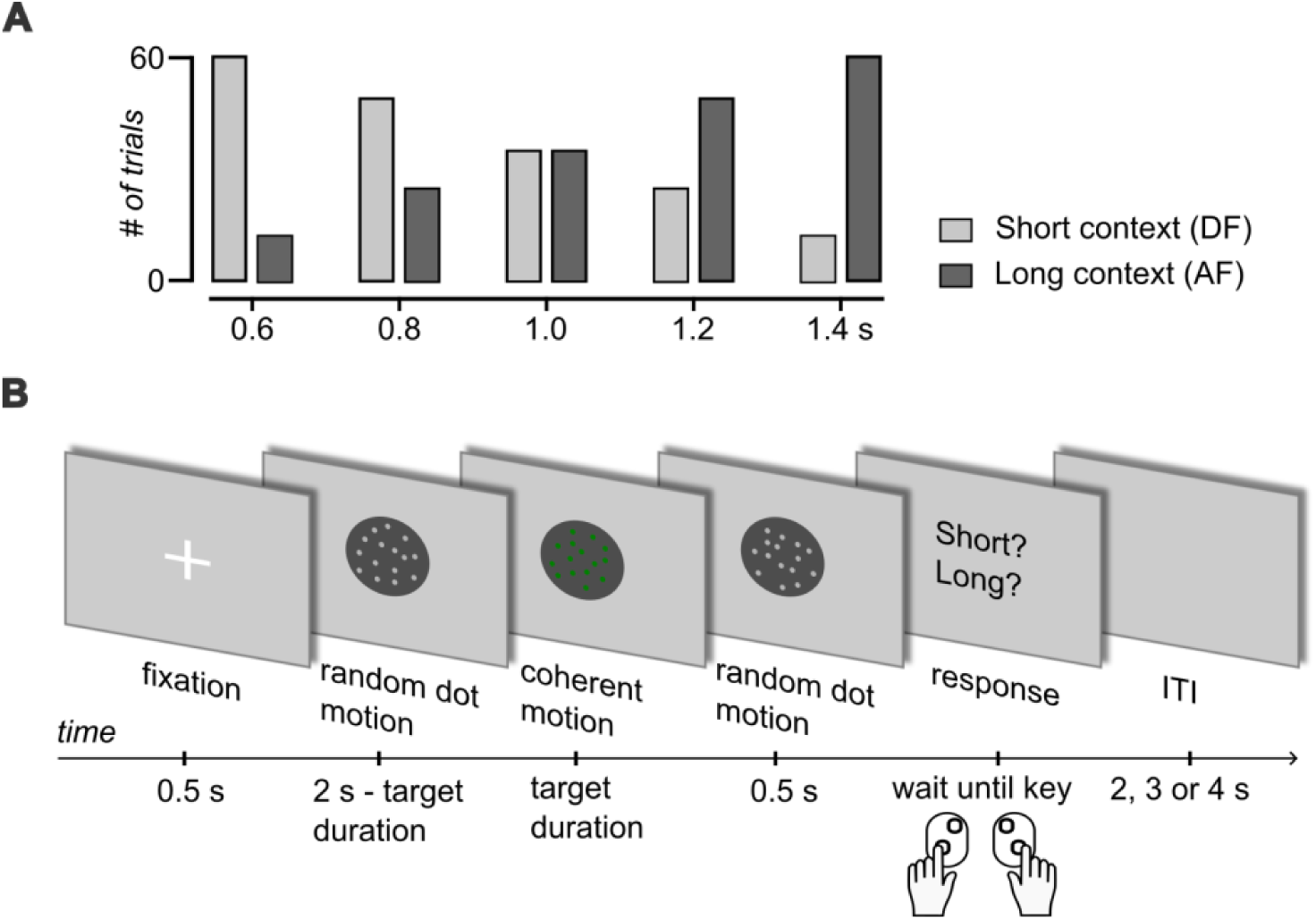
(A) Target intervals and context distributions. Five durations (0.6, 0.8, 1.0, 1.2, and 1.4 s) were presented with different frequencies across conditions. In the descending-frequency (DF) context, short durations were more frequent, whereas in the ascending-frequency (AF) context, long durations were more frequent. **(B)** Trial structure: fixation, random-dot motion, coherent target interval, response prompt, and variable ITI.

Stimuli were projected onto a rear-projection canvas (diagonal 30 in.) by an MRI-compatible ProPIXX DLP projector (Pixx Technologies Inc., Canada) and viewed through a head-coil-mounted mirror at 110 cm. The display was a random-dot kinematogram (15 white dots, diameter 0.32°, speed 7°/s) inside an 11.6°-diameter dark-gray circular aperture (RGB [64, 64, 64]) on a light-gray background (RGB [128, 128, 128]).

Each trial (Figure 1B) had a 0.5-s fixation cross, a 2.5-s moving-dot display (random motion-white color for 2 s − target duration, coherent motion-green color for the target duration, then random motion-white color for 0.5 s), a “Short or Long?” response prompt, and an inter-trial interval (ITI) of 2, 3, or 4 s. The total 2.5-s display duration was fixed across target durations and contexts to equate visual stimulation and run-onset timing; coherent-motion direction was held constant (0°, rightward); the short/long key mapping was counterbalanced across participants.

Before the main experiment, participants completed 5–10 practice trials with feedback using only the anchor durations (0.6 and 1.4 s), outside the scanner. The main experiment used five equally spaced target durations (0.6, 0.8, 1.0, 1.2, 1.4 s) presented at unequal frequencies across two context conditions (Figure 1A): in the descending-frequency (DF) context the short to long counts were 60/48/36/24/12 trials, and in the ascending-frequency (AF) context they were 12/24/36/48/60 trials. AF and DF were tested in separate functional runs (180 trials per run, 360 total), each comprising six 30-trial blocks separated by a 6-s rest. Within every run, the five target durations were interleaved according to a per-subject sequence generated offline (no fixed within-block order). Context-run order was randomly assigned across participants (13 participants, first AF then DF, 11 participants, first DF then AF), and we revisit run-order as a sensitivity covariate below. Between functional runs, structural imaging was acquired (∼6 minutes).

#### Paradigm considerations

We replaced the auditory tones used in our group’s earlier bisection work (Zhu et al., 2021; Baykan et al., 2023) with a visual moving-dot stimulus because random-dot kinematograms are MRI-compatible, avoid scanner-noise masking, and allow tight control over stimulus presentation. Two design features bear on the analyses below. First, because the coherent target always occupied the final segment of the fixed 2.5-s display, its offset was constant and its onset latency was the complement of its duration (the random-motion lead-in lasted *2.0 s − Target Duration*). The display therefore provides two redundant cues to the same duration judgment: (i) the duration of the coherent target, and (ii) its onset latency, so in principle participants could use both cues for the bisection task, although the instruction emphasized judgments based on the coherent motion. As a cautious approach, we accordingly interpret duration-related signals as duration-related but cue-ambiguous, and analyse the data under two duration-modulator alignments, at trial onset and at coherent-target onset (see *First-level general linear models*). Second, visual motion can itself bias perceived duration (Kanai et al., 2006; Kaneko & Murakami, 2009); however, because direction and speed were constant across contexts, motion cannot explain AF–DF differences but warrants cautious phrasing of absolute duration claims as duration-related rather than pure-duration encoding.

### Behavioral Analysis

For each participant and condition we fitted a cumulative-Gaussian psychometric function with a symmetric lapse parameter, *P*(long | *x*) = λ + (1 − 2λ) · Φ((*x* − μ) / σ), by binomial maximum likelihood, where μ is the point of subjective equality (PSE), σ is the dispersion, and λ is the per-side lapse rate. We report the λ ≤ 0.10 fits as the headline psychometric model (few fits at the upper bound) and λ ≤ 0.05 as a stricter sensitivity check, alongside the original lapse-free fits. The Weber fraction (σ / μ) indexed temporal precision.

The context effect on accuracy was tested as the PSE shift (PSE_AF_ − PSE_DF_) with a paired *t*-test. For precision we report (i) a frequentist paired *t*-test on the AF–DF Weber-fraction difference, (ii) a Bayesian paired *t*-test (Rouder et al., 2009) with prior-scale sensitivity (*r* = 0.5, 0.707, 1.0), and (iii) a two one-sided tests (TOST) equivalence procedure (Lakens, 2017) against the auditory-bisection effect size of Zhu et al. (2021), so that a null is interpreted as evidence for equivalence rather than as a non-significant difference.

### Image Acquisition

Data were acquired on a 3T Siemens MAGNETOM Prisma scanner with a 32-channel head coil. Functional images used a T2*-weighted multiband EPI sequence (TR = 1000 ms, TE = 30 ms, flip angle = 45°, FOV = 210 × 210 mm², 3-mm isotropic voxels, 48 axial slices, multiband factor = 4). T1-weighted structural images were acquired with an MPRAGE sequence (TR = 2500 ms, TE = 2.22 ms, flip angle = 8°, FOV = 256 × 256 mm², 0.8-mm isotropic voxels).

### Image Preprocessing

Preprocessing was performed using *fMRIPrep* 23.1.4 (Esteban et al., 2020). In brief, structural images were skull-stripped, segmented, and normalized to MNI152NLin2009cAsym space. BOLD data underwent motion correction, slice-timing correction, co-registration to the structural image, and resampling to MNI space (2 mm isotropic). All 24 participants met the a priori motion criterion (mean FD < 0.5 mm) and were retained. The full fMRIPrep boilerplate description is provided in Appendix A.

### Statistical Analysis

#### First-Level General Linear Models

First-level analyses used *nilearn* (Huntenburg et al., 2017) on data spatially smoothed at 6-mm FWHM. Each participant’s model contained, per context, a 2.5-s constant-amplitude boxcar at trial onset capturing the duration-invariant trial response (AF, DF) and a mean-centred parametric modulator (Target Duration − 1.0 s) carrying duration-related modulation (AF_Dur_, DF_Dur_). Mean-centering reduced collinearity between the duration modulator and the context boxcar.

We estimated two first-level GLMs that differ only in where the duration modulator is placed in time. In the *trial-onset GLM*, the modulator was a delta function at trial onset. In the *target-onset GLM*, the modulator was re-time-locked to the onset of the coherent target, and a response-prompt stick regressor was added at *trial onset + 2.5 s* as a coarse motor-control regressor (trial-wise reaction times were not retained in this dataset’s BIDS events, only the choice code, 1or 2; see Appendix C). Because the coherent target, not the trial onset, is the event whose duration is modulated, and because the target-onset GLM additionally models the response, we foreground the target-onset GLM as the analysis of record and report the trial-onset GLM throughout as a comparison. The two alignments are not interchangeable: the target’s duration and its onset latency are complementary cues by the fixed display window (see *Paradigm considerations*), so the placement of the modulator is itself consequential. Decision rules for comparing the two GLMs were fixed before the target-onset GLM was estimated and are detailed in Appendix B.

All task regressors were convolved with the canonical Glover HRF. Nuisance regressors comprised six rigid-body motion parameters and six aCompCor components. We high-pass-filtered at 1/128 Hz (discrete cosine basis) and modeled temporal autocorrelation with AR(1).

First-level contrasts were (1) task activation, (AF + DF) vs. baseline; (2) context main effect, AF − DF; (3) collapsed duration encoding, AF_Dur_ + DF_Dur_; (4) context-specific duration encoding, AF_Dur_ and DF_Dur_; (5) context × duration interaction, AF_Dur_ − DF_Dur_.

#### Design-matrix diagnostics

For both the trial-onset and target-onset GLMs we exported per-subject, per-run pairwise Pearson correlations among available task regressors and the VIF of each task regressor after regressing it on all other non-constant design columns. Full diagnostic tables appear in Appendix D.

#### Second-Level Analysis

Group-level random-effects analyses used one-sample *t*-tests on first-level contrast estimates. Whole-brain maps were thresholded at *z* > 3.1 (*k* > 10 voxels) with permutation-based FWE correction (5000 permutations).

#### Regions of Interest and Small-Volume Correction

From activation-likelihood-estimation meta-analyses of temporal processing (Mondok & Wiener, 2022), we defined a priori 8-mm-radius spherical ROIs at the following MNI coordinates (Figure 2): SMA (0, 12, 50; Wiener et al., 2010), bilateral putamen (±24, 4, 2), bilateral insula (±38, 14, 0), bilateral cerebellum lobule VI (±28, −60, −26), and bilateral IFG (±52, 18, 8) (Wiener et al., 2010; Nani et al., 2019; Wittmann et al., 2010; Coull et al., 2004). Small-volume correction was applied within each sphere using voxel-level Bonferroni correction (two-tailed; *p*FWE-SVC < .05); we additionally report a strict cross-ROI Bonferroni threshold (*p* < .00556) as a conservative benchmark.

**Figure 2.**
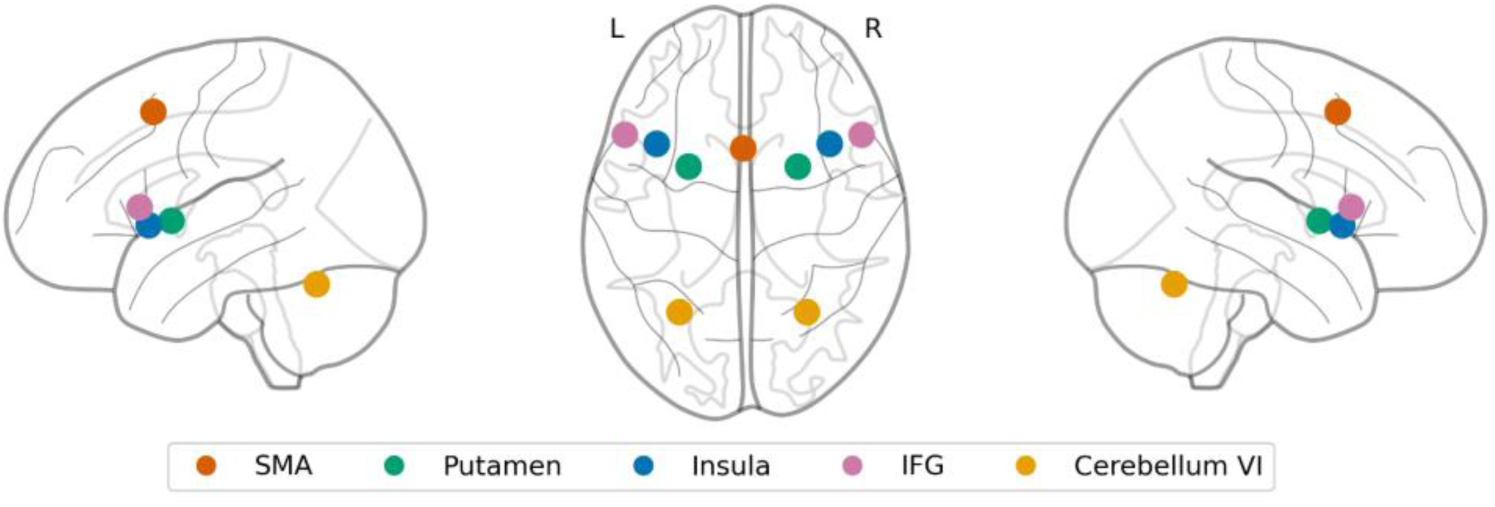
A priori regions of interest for temporal processing. Glass brain projections (left sagittal, axial, right sagittal views) show the nine ROI spheres (8 mm radius) centered on coordinates from the neuroimaging timing literature: SMA (red), bilateral putamen (green), bilateral insula (blue), IFG (purple), and bilateral cerebellum lobule VI (orange).

### ROI Dissociation: Context × Region Interaction

To test whether duration-related activity was differentially distributed across the timing network, we extracted mean contrast estimates from AF_Dur_ and DF_Dur_ in each a priori ROI (bilateral ROIs averaged across hemispheres) and submitted them to a 2 (Context) × 5 (Region: SMA, putamen, insula, cerebellum VI, IFG) repeated-measures ANOVA with Greenhouse–Geisser correction (reported as *F*, *p*_GG_, partial η²). AF vs. DF at each of the five a priori ROIs was then tested with planned paired *t*-tests, Holm-corrected across the five tests; lateralized (per-hemisphere) follow-ups are retained in Appendix D as descriptive analyses. The 2 × 5 ANOVA and Holm-corrected simple effects were computed for the foregrounded target-onset GLM and, for comparison, the trial-onset GLM; decision rules for this comparison were fixed before the target-onset GLM was estimated (Appendix B). We additionally added run-order (AF-first vs. DF-first) as a between-subjects covariate.

#### Generalized Psychophysiological Interaction (gPPI) Analysis

We probed whether SMA functional coupling during duration encoding varied with temporal context using a generalized psychophysiological interaction analysis (McLaren et al., 2012) implemented in custom Python. The seed was an 8-mm sphere centred on the SMA AF_Dur_ − DF_Dur_ peak (MNI 6, −4, 58); the *a priori* SMA ROI sphere (0, 12, 50) was used as a seed-sensitivity check. For each participant and run we extracted the seed BOLD time series, recovered the underlying neural signal by Wiener deconvolution of the canonical HRF, multiplied it by each psychological variable, and reconvolved with the HRF. We computed PPI contrasts for the parametric interaction difference (PPI_AFT_ − PPI_DFT_) and the non-parametric interaction difference (PPI_AF_ − PPI_DF_), thresholded at *z* > 3.1, *k* > 10 voxels, and report each contrast in both directions (AF > DF, DF > AF). Per-subject diagnostics of the parametric and non-parametric PPI interaction terms (L2 norm, temporal variance, parametric–non-parametric correlation) were computed so that any detectability asymmetry could be attributed to regressor structure rather than neural connectivity; across subjects, parametric–non-parametric correlations were modest in both seeds (max |*r*| = 0.32, below the 0.5 group-level threshold specified in Appendix B), with one subject (*a priori*, sub-020 variance outlier with clean *r* values) flagged but no subject excluded. The gPPI is treated as exploratory; whole-brain cluster tables appear in Appendix E.

#### Brain–Behavior Correlation

To test whether individual neural context modulation tracked behavioral sensitivity, we correlated each participant’s AF_Dur_ − DF_Dur_ contrast estimates (extracted from 8-mm spheres at SMA, right putamen, right IFG, and left insula) with the PSE shift (PSE_AF_ − PSE_DF_), using Pearson correlations and partial correlations controlling for the Weber fraction.

## Results

### Behavioral Results

Temporal context produced a robust shift in subjective duration judgments. Cumulative-Gaussian psychometric fits with a symmetric bounded lapse parameter (λ ≤ .10; headline model) yielded mean PSEs of 1.041 s (*SD* = 0.129) in the AF context and 0.890 s (*SD* = 0.093) in the DF context, a 151 ms shift (95% CI [0.106, 0.195]), *t*(23) = 6.93, *p* < .001 (Figure 3B). A stricter lapse bound (λ ≤ .05) and an unbounded-lapse fit with λ freely estimated returned the same results (Appendix C).

**Figure 3.**
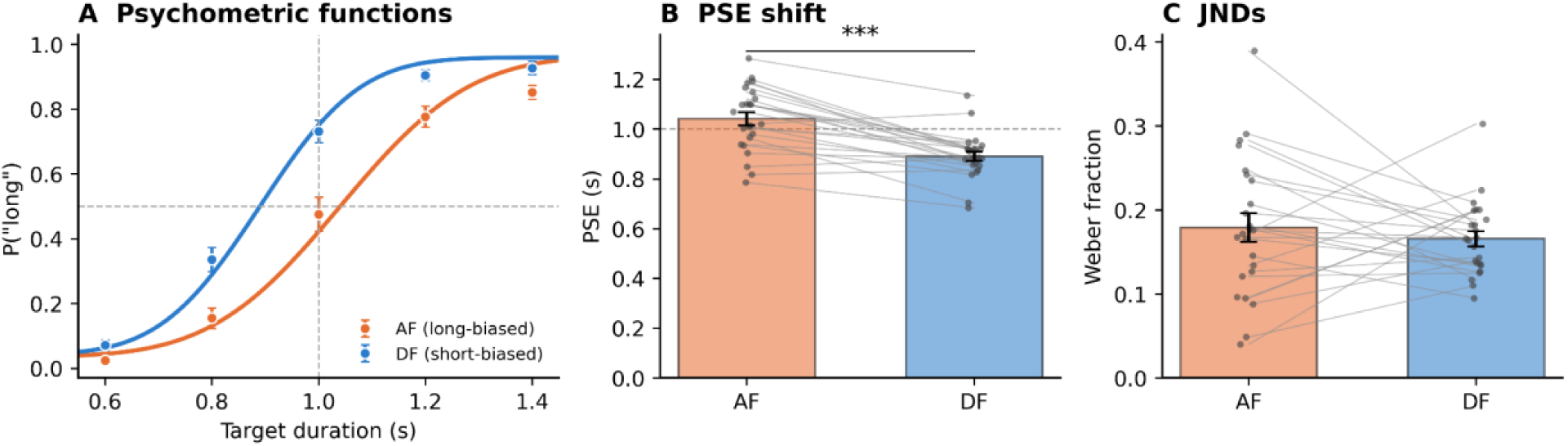
Behavioral context effect in the duration bisection task. **(A)** Lapse-corrected cumulative-Gaussian fits for AF (long-biased) and DF (short-biased) contexts with group mean response proportions. **(B)** PSE values by context with individual participants and group means. **(C)** Weber fractions by context; annotation summarizes Bayesian and equivalence-test evidence.

Temporal precision was comparable across contexts. Weber fractions averaged 0.179 (AF) and 0.166 (DF), Δ = 0.013, *t*(23) = 0.72, *p* = .476, *d* = 0.15. A JZS paired *t*-test returned BF_01_ = 3.67, indicating moderate evidence for no AF–DF Weber-fraction difference. The TOST equivalence test supported equivalence against a liberal ±0.06 Weber-unit bound (*p*TOST = .010), but not against the stricter ±0.04 bound (*p*_TOST_ = .083). Thus, context shifted the decision criterion without reliable evidence for a precision change. The PSE shift was already present in early blocks (153 ms) and did not increase in late blocks (143 ms), Context × Block Position *p* = .742, suggesting rapid adaptation to the distributional context.

### fMRI Results

#### Engagement of the core timing network

The Task contrast (AF + DF vs. baseline) recruited the expected timing network, with widespread bilateral activation spanning SMA, premotor cortex, IFG, inferior parietal lobule, basal ganglia, insula, and cerebellum (12,199 voxels at *z* > 3.1, peak *z* = 6.45; all clusters *p*_FWE_ < .05, whole-brain cluster-level corrected).

Collapsed across contexts (AF_Dur_ + DF_Dur_), the target-onset GLM showed duration-related modulation reaching small-volume significance in the right IFG (peak *z* = 3.47, *p*_FWE-SVC_ = .039, MNI: 56, 18, 7), with a trend in SMA/pre-SMA (peak *z* = 3.27, *p* = .079); no other a priori ROI approached significance.

#### Context-specific duration modulation

We foreground the target-onset GLM, which time-locks the duration modulator to the coherent target and includes an explicit response regressor (Methods, *First-level general linear models*); the registered trial-onset GLM is reported throughout as a comparison. Small-volume correction within the a priori ROIs localized duration-related modulation predominantly to the short-biased (DF) context (Table 1). In the DF context, duration-related modulation survived SVC in SMA/pre-SMA (peak *z* = 3.73, *p*_FWE-SVC_ = .014, MNI: 0, 6, 53), the right insula (peak *z* = 3.73, *p* = .014, MNI: 42, 12, 4), and the right IFG (peak *z* = 3.50, *p* = .035), with trends in the left insula and right cerebellum. In the AF context, no a priori ROI survived correction. The Context × Duration contrast (AF_Dur_ − DF_Dur_) confirmed this asymmetry: the right insula showed a significant DF > AF interaction (peak *z* = −3.49, *p*_FWE-SVC_ = .035, MNI: 42, 14, 4; Figure 4), the only a priori ROI to survive small-volume correction.

**Figure 4.**
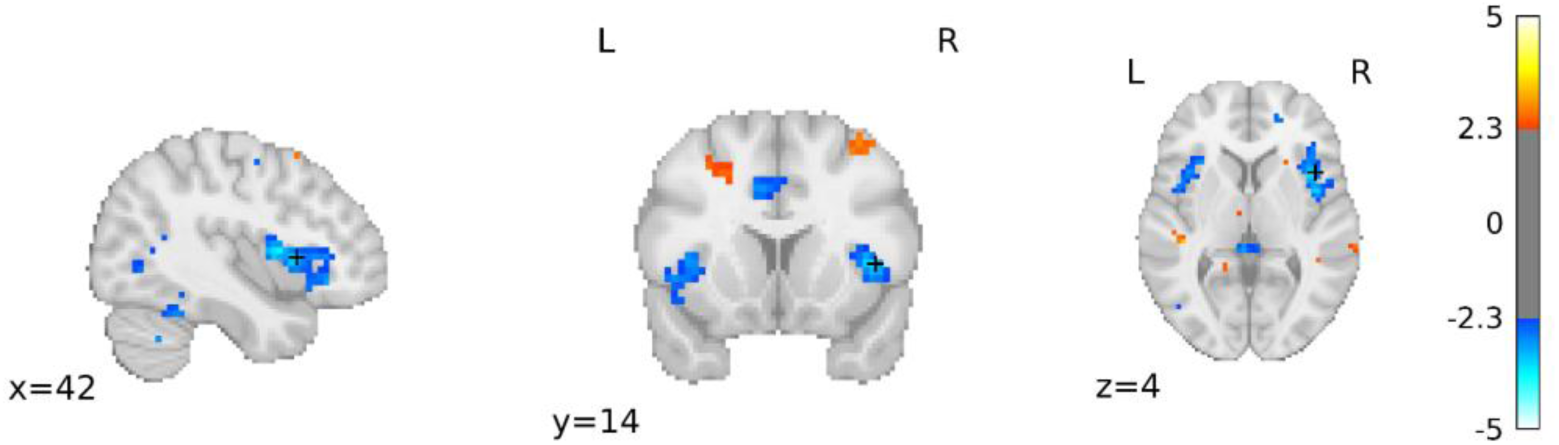
Whole-brain context-by-duration map (AF_Dur_ − DF_Dur_) from the target-onset GLM. Warm colors indicate stronger AF than DF duration-related modulation; cool colors indicate the reverse (DF > AF). Maps are displayed at *z* > 2.3; the crosshair marks the a priori right-insula peak (MNI 42, 14, 4), the only ROI whose Context × Duration peak survives small-volume correction.

**Figure 5.**
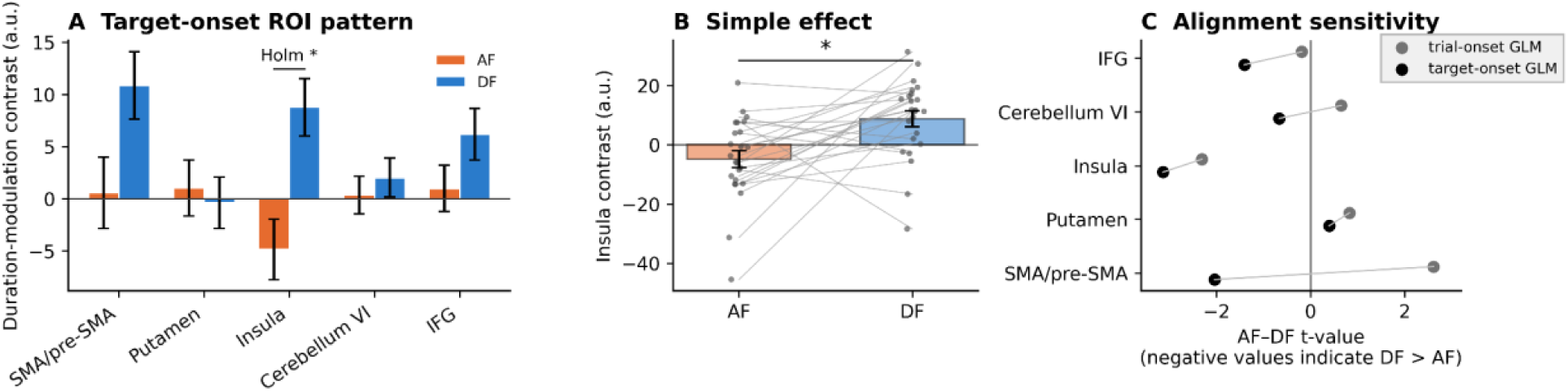
ROI dissociation (target-onset GLM). **(A)** Target-onset GLM contrast estimates across the five bilateral ROI families. **(B)** Left-insula simple effect, the only Holm-corrected ROI simple effect. **(C)** Trial-onset versus target-onset simple-effect *t*-values, illustrating that the SMA simple effect is alignment-sensitive — reversing sign between alignments — whereas the insula DF > AF effect is consistent and strengthens under target-onset alignment.

**Table 1.** SVC peak z-statistics (FWE-corrected p-values) within 8-mm spherical a priori ROIs for key contrasts from the target-onset GLM (the foregrounded analysis). Bold indicates *p*FWE-SVC < .05. Values are the signed peak z within each search volume; for Context × Duration, positive denotes AF > DF and negative DF > AF. The corresponding trial-onset GLM table is given in Appendix D.

| ROI | Duration $z$ ($p$) | $AF_{Dur}$ $z$ ($p$) | $DF_{Dur}$ $z$ ($p$) | Context × |
| --- | --- | --- | --- | --- |
| | | | | Duration $z$ ($p$) |
| SMA | 3.27 (.079) | 1.44 (1.00) | **3.73 (.014)** | **-2.71 (.505)** |
| R Putamen | 2.24 (1.00) | 2.89 (.261) | 1.08 (1.00) | 2.07 (1.00) |
| L Putamen | 1.23 (1.00) | -1.65 (1.00) | -1.62 (1.00) | 1.91 (1.00) |
| R IFG | **3.47 (.039)** | 3.19 (.106) | **3.50 (.035)** | -2.26 (1.00) |
| L IFG | 2.09 (1.00) | -1.59 (1.00) | 2.35 (1.00) | -1.85 (1.00) |
| R Insula | 2.04 (1.00) | -3.32 (.064) | **3.73 (.014)** | **-3.49 (.035)** |
| L Insula | 2.03 (1.00) | -2.29 (1.00) | 3.24 (.088) | -2.92 (.262) |

| ROI | Duration $z$ ($p$) | AF<sub>Dur</sub> $z$ ($p$) | DF<sub>Dur</sub> $z$ ($p$) | Context × |
| --- | --- | --- | --- | --- |
| | | | | Duration $z$ ($p$) |
| R Cerebellum VI | 2.68 (.540) | 1.60 (1.00) | 3.31 (.069) | -2.54 (.816) |
| L Cerebellum VI | 1.58 (1.00) | 1.43 (1.00) | 1.39 (1.00) | -1.88 (1.00) |

This regional pattern is specific to the target-onset alignment. Under the trial-onset GLM, duration-related modulation instead loaded on the AF context, surviving SVC in SMA/pre-SMA, right putamen, and right IFG, and the Context × Duration interaction peaked in SMA/pre-SMA in the AF > DF direction (*z* = 3.45, *p*FWE-SVC = .041; full trial-onset SVC table in Appendix D). Because the coherent target’s duration and its onset latency are complementary cues by the fixed display window (Methods, *Paradigm considerations*), the regional attribution of duration-related signal, and the sign of the SMA interaction, shifts with where the duration modulator is placed. We treat this alignment-sensitivity as a substantive limitation and return to it in the Discussion.

#### Regional specificity and robustness

A 2 (Context) × 5 (Region) repeated-measures ANOVA on bilateral mean contrast estimates tested whether distributional context changed the regional distribution of duration-related activity. In the target-onset GLM, the Context × Region interaction was significant, *F*(4, 92) = 3.97, *p*_GG_ = .014, 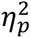 = .147, supporting a network-level context effect. Planned AF-vs-DF paired *t*-tests at each ROI, Holm-corrected across the five tests, showed that the left-insula DF > AF effect survived correction, *t*(23) = −3.13, *p*Holm = .023; no other ROI simple effect was reliable (Table 2).

**Table 2.**
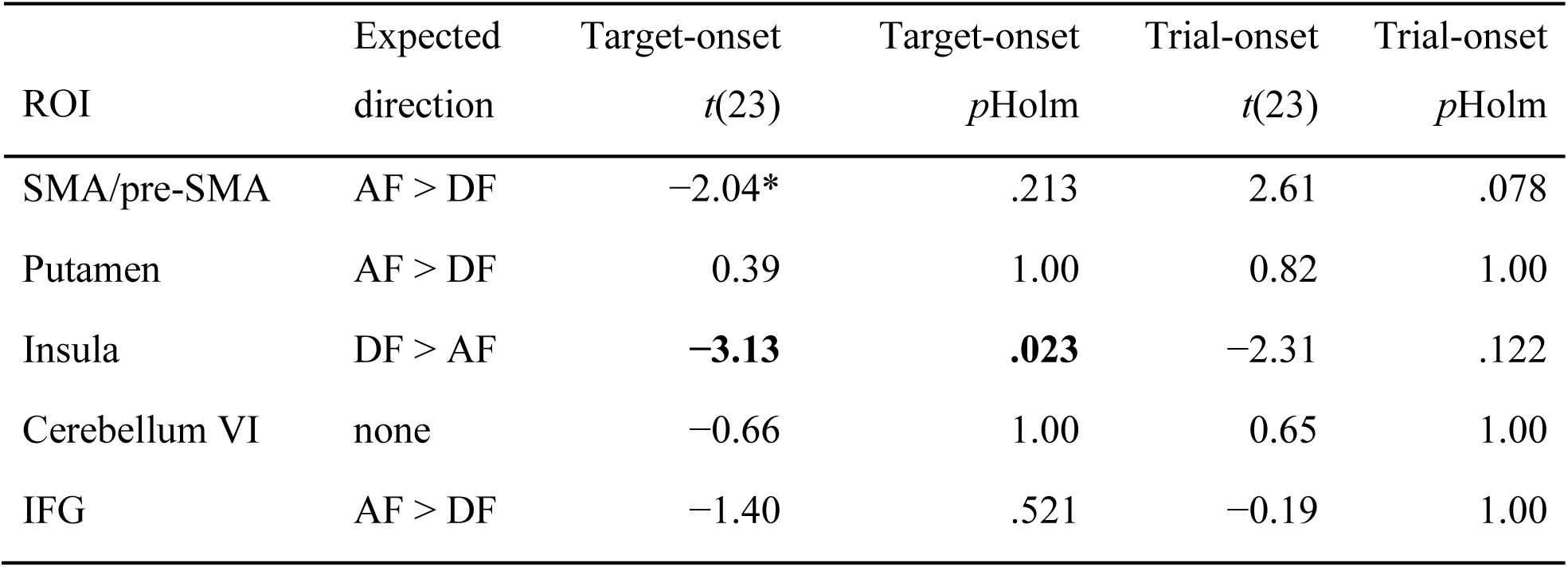
Planned AF-vs-DF paired tests at each a priori ROI, Holm-corrected across the five tests, shown for the target-onset GLM (foregrounded) and the trial-onset GLM (comparison). ROIs are bilateral averages. Bold indicates a Holm-corrected effect at α = .05; the asterisk marks the SMA sign reversal between alignments.

The registered trial-onset GLM returned the same network-level conclusion: the Context × Region interaction was significant, *F*(4, 92) = 5.90, *p*_GG_ = .001, 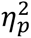 = .204. Its simple-effect profile differed, however, no ROI survived Holm correction, and the SMA effect ran in the AF > DF direction, opposite to the target-onset GLM (Table 2), the same alignment-sensitivity seen in the whole-brain SVC. Adding run-order (AF-first vs. DF-first) as a between-subject covariate did not change the conclusion: the Context × Region interaction remained significant under both alignments, and the Block-order × Context × Region interaction was non-significant.

Together, the ROI analyses support a network-level Context × Region effect, with the left insula providing the clearest and most alignment-stable region-specific evidence. SMA/pre-SMA contributes to the network-level pattern, but its duration-related simple effect reverses sign between the two regressor alignments and is therefore not treated as a robust region-specific claim.

#### Context-dependent SMA functional coupling and brain–behavior association

Exploratory generalized PPI analyses indicated that SMA-centered functional coupling during duration-related processing varied with context. The parametric AF_Dur_ > DF_Dur_ and DF_Dur_ > AF_Dur_ contrasts converged across the data-driven and a priori SMA seeds on the same peak partners (right IFG/pre-motor cortex and right V5/MT+). The non-parametric context PPI contrasts produced no clusters in either seed, consistent with the fact that the parametric PPI term carried trial-by-trial duration information whereas the boxcar context term did not. No a priori ROI-to-ROI coupling contrast survived FDR correction, and full cluster and diagnostic tables are reported in Appendix E.

Individual differences in neural context modulation did not track behavioral PSE shifts. Pearson correlations between AF_Dur_ − DF_Dur_ contrast estimates at SMA, right putamen, right IFG, and left insula and the AF − DF PSE shift were all non-significant; partial correlations controlling for Weber fraction returned the same pattern. These analyses provide no evidence for a reliable brain–behavior association, but the sample remains underpowered for small individual-difference effects.

## Discussion

This present study examined how distributional context changes neural processing during visual temporal bisection. Behaviorally, long- and short-biased duration distributions shifted subjective duration judgments. The results replicate the established ensemble-distribution effect: recent temporal statistics biased the bisection criterion. This pattern fits Bayesian and ensemble-distribution accounts in which observers integrate current sensory evidence with a contextual prior derived from recent samples (Jazayeri & Shadlen, 2010; Zhu et al., 2021). Neurally, context modulated duration-related activity across an established timing network. The strongest evidence is a network-level Context × Region pattern across a priori timing ROIs, with the left insula the clearest region-specific locus. SMA/pre-SMA remains theoretically important, but its duration-related simple effect reversed sign with duration-regressor alignment and is therefore best interpreted at the network level. Our findings support context-sensitive weighting within timing-related circuitry rather than recruitment of anatomically separate timing systems.

The main contribution of the present fMRI study is spatial and mechanistic. Prior behavioral and EEG work established that distributional context shifts temporal decisions and modulates temporally extended neural dynamics, including anticipatory and post-stimulus components (Baykan et al., 2023). The present data show that this contextual influence is expressed across a priori timing-related regions, rather than being confined to a single dedicated region. In this sense, the study extends the EEG findings by showing that this contextual influence is distributed across the timing network, most clearly expressed in the insula.

### Context-dependent weighting across the timing network

The left-insula result provides the clearest region-specific evidence. Insular cortex has been associated with interoceptive timing, salience, and subjective duration (Craig, 2009; Wittmann, van Wassenhove, et al., 2010). Stronger duration-related modulation in the short-biased context may reflect increased weighting of bodily, salience-related, or decision-related timing signals when shorter intervals are more diagnostic. This should not be read as evidence that the insula is a separate short-duration timing system. Rather, it indicates that the relative contribution or detectability of insular duration-related signals changed with distributional context.

The right IFG showed duration-related modulation both overall and within the short-biased context, consistent with fronto-cortical accounts of temporal attention and accumulation (Matell & Meck, 2004; Wiener et al., 2010); putaminal and cerebellar effects were weaker and alignment-dependent, where the cerebellar duration sensitivity present under trial-onset alignment did not survive target-onset alignment. Several non-mutually exclusive mechanisms could account for this context-dependent redistribution of duration-related modulation. A predictive-coding account frames it as reweighting of duration-related signals relative to a distributional prior, through the precision afforded to prediction error (Jazayeri & Shadlen, 2010; Shi et al., 2013); a fronto-cortical attention account links the IFG involvement to context-dependent allocation of temporal attention (Coull et al., 2004). Both predict that context reweights duration-related activity, but neither uniquely predicts its direction in neural activities. The concentration in the short-biased context is most naturally accommodated by an interoceptive or salience account, on which insular contributions grow when shorter intervals or categorical boundaries become behaviorally diagnostic (Craig, 2009; Wittmann, Simmons, et al., 2010). These accounts converge on differential weighting within a shared timing network, and may in part reflect greater detectability of duration-related neural signals in that context (as noted above) rather than a mechanistic asymmetry.

### SMA/pre-SMA and context-sensitive timing

SMA/pre-SMA remains a plausible site for contextual modulation because medial frontal timing regions have been linked to temporal accumulation, expectation, and prior-guided decision boundaries (Cui et al., 2009; Mita et al., 2009). Ramping activity in SMA/pre-SMA is often interpreted as supporting temporal prediction or motor simulation (Merchant et al., 2013), and SMA activity scales with both objective and subjectively perceived duration (Coull et al., 2016). Causal evidence also converges on this region: stimulation of SMA can alter temporal bisection precision or bias (Méndez et al., 2017; Wiener et al., 2018), and dorsal premotor cortex has been implicated in temporal prediction (Lazzari et al., 2025).

The present findings are consistent with this literature, but they refine an SMA-centered interpretation. SMA/pre-SMA contributed to the network-level Context × Region effect, and in the foregrounded target-onset GLM showed duration-related modulation in the short-biased (DF) context. Its Context × Duration *simple effect*, however, reversed sign between the two duration-modulator alignments: AF > DF when the modulator was placed at trial onset, DF > AF when it was placed at coherent-target onset. This alignment-sensitivity is expected given the task structure: because the coherent target had a fixed offset within the 2.5-s display, its duration and its onset latency were complementary, perfectly anti-correlated cues, and a duration-related signal can be carried by either. Moving the modulator between trial onset and target onset re-weights these two cues and, because they are anti-correlated, can reverse the sign of the estimated interaction. The reversal is consistent with SMA/pre-SMA tracking *when* the coherent target appears, an onset-latency or anticipatory, hazard-like timing signal, as much as its duration, but the design cannot identify which cue drives the signal. The safer interpretation is that SMA/pre-SMA participates in context-sensitive timing, while the present design does not isolate a robust SMA-specific duration effect.

This interpretation is compatible with Bayesian accounts of temporal perception. SMA activity has been linked to cumulative hazard derived from the prior distribution rather than elapsed time alone (Cui et al., 2009), and primate pre-SMA recordings show that prior distributions can warp neural dynamics in a manner consistent with Bayesian integration (Sohn et al., 2019). The current fMRI data place SMA/pre-SMA within this context-sensitive timing network, but they do not require it to be the sole or most robust locus of distributional-context effects.

### Exploratory connectivity, limitations, and future directions

The SMA-seeded gPPI results provide exploratory converging evidence that context modulated duration-related coupling with timing- and attention-related regions. Because the parametric and non-parametric PPI terms differ in temporal structure, the absence of non-parametric clusters should not be interpreted as definitive evidence for duration-specific connectivity. A more cautious conclusion is that SMA-centered coupling is sensitive to distributional context in a manner consistent with the regional analyses, but not independently confirmatory of a distinct connectivity mechanism.

Several limitations constrain interpretation. First, the coherent target had a fixed offset within the 2.5-s display, so its duration and its onset latency were complementary, perfectly anti-correlated cues to the bisection judgment. The design, therefore, cannot establish whether duration-related modulation reflects interval timing of the target, anticipation of its onset, or both. We foreground the target-onset GLM because it places the duration modulator at the event whose duration is modulated and because it models the response; but re-aligning the modulator does not, and cannot, separate cues that are confounded by construction. Therefore, the regional attribution of duration-related signal, and the sign of the SMA interaction, differ between the two alignments. Addressing this requires a new event-related design that decouples target onset from target duration. Second, the blocked design stabilized distributional context but cannot resolve trial-by-trial updating. Third, the linear parametric modulator tests monotonic duration sensitivity and may miss duration-tuned responses. Finally, the gPPI and brain–behavior analyses should be treated as exploratory given regressor-structure constraints and limited power for individual differences.

In summary, distributional context biased temporal bisection judgments and modulated duration-related activity across established timing regions. The evidence supports a network-level account: context changes the relative weighting of timing-related signals, with the left insula providing the clearest region-specific evidence and SMA/pre-SMA contributing to the broader timing-network pattern. These findings extend behavioral and EEG evidence for temporal-context effects by identifying where such effects are expressed in human fMRI. Future event-related designs that decouple target duration from temporal expectation, combined with nonlinear duration models, will be needed to distinguish duration encoding, prior weighting, and expectation-related signals within this network.

