## Supplementary Materials for "Distributional context reweights duration-related activity across timing-related regions"

### Appendix A: fMRIPrep Preprocessing Details

The following boilerplate text was automatically generated by fMRIPrep with the express intention that users should copy and paste this text into their manuscripts unchanged. It is released under the CC0 license.

Results included in this manuscript come from preprocessing performed using *fMRIPrep* 23.1.4 (Esteban et al., 2019; Esteban et al., 2020; RRID:SCR_016216), which is based on *Nipype* 1.8.6 (Gorgolewski et al., 2011; Gorgolewski et al., 2018; RRID:SCR_002502).

#### Anatomical Data Preprocessing

A total of 1 T1-weighted (T1w) image was found within the input BIDS dataset. The T1-weighted (T1w) image was corrected for intensity non-uniformity (INU) with N4BiasFieldCorrection (Tustison et al., 2010), distributed with ANTs, and used as T1w-reference throughout the workflow. The T1w-reference was then skull-stripped with a *Nipype* implementation of the antsBrainExtraction.sh workflow (from ANTs), using OASIS30ANTs as target template. Brain tissue segmentation of cerebrospinal fluid (CSF), white-matter (WM) and gray-matter (GM) was performed on the brain-extracted T1w using fast (FSL, RRID:SCR_002823; Zhang et al., 2001). Volume-based spatial normalization to one standard space (MNI152NLin2009cAsym) was performed through nonlinear registration with antsRegistration (ANTs), using brain-extracted versions of both T1w reference and the T1w template. The following template was selected for spatial normalization and accessed with *TemplateFlow* (23.0.0; Ciric et al., 2022): *ICBM 152 Nonlinear Asymmetrical template version 2009c* (Fonov et al., 2011; RRID:SCR_008796; TemplateFlow ID: MNI152NLin2009cAsym).

#### Functional Data Preprocessing

For each of the 2 BOLD runs found per subject (across all tasks and sessions), the following preprocessing was performed. First, a reference volume and its skull-stripped version were generated using a custom methodology of *fMRIPrep*. Head-motion parameters with respect to the BOLD reference (transformation matrices, and six corresponding rotation and translation parameters) were estimated before any spatiotemporal filtering using mcflirt (FSL; Jenkinson et al., 2002). BOLD runs were slice-time corrected to 0.449s (0.5 of slice acquisition range 0s–0.897s) using 3dTshift from AFNI (Cox & Hyde, 1997; RRID:SCR_005927).

The BOLD time-series (including slice-timing correction when applied) were resampled onto their original, native space by applying the transforms to correct for head-motion. These resampled BOLD time-series will be referred to as *preprocessed BOLD in original space*, or just *preprocessed BOLD*. The BOLD reference was then co-registered to the T1w reference using mri_coreg (FreeSurfer) followed by flirt (FSL; Jenkinson & Smith, 2001) with the boundary-based registration (Greve & Fischl, 2009) cost-function. Co-registration was configured with six degrees of freedom.

Several confounding time-series were calculated based on the *preprocessed BOLD*: framewise displacement (FD), DVARS and three region-wise global signals. FD was computed using two formulations following Power (absolute sum of relative motions; Power et al., 2012) and Jenkinson (relative root mean square displacement between affines; Jenkinson et al., 2002). FD and DVARS are calculated for each functional run, both using their implementations in *Nipype* (following the definitions by Power et al., 2012). The three global signals are extracted within the CSF, the WM, and the whole-brain masks.

Additionally, a set of physiological regressors were extracted to allow for component-based noise correction (*CompCor*; Behzadi et al., 2007). Principal components are estimated after high-pass filtering the *preprocessed BOLD* time-series (using a discrete cosine filter with 128s cut-off) for the two *CompCor* variants: temporal (tCompCor) and anatomical (aCompCor). tCompCor components are then calculated from the top 2% variable voxels within the brain mask. For aCompCor, three probabilistic masks (CSF, WM and combined CSF+WM) are generated in anatomical space. The implementation differs from that of Behzadi et al. in that instead of eroding the masks by 2 pixels on BOLD space, a mask of pixels that likely contain a volume fraction of GM is subtracted from the aCompCor masks. This mask is obtained by thresholding the corresponding partial volume map at 0.05, and it ensures components are not extracted from voxels containing a minimal fraction of GM. Finally, these masks are resampled into BOLD space and binarized by thresholding at 0.99 (as in the original implementation). Components are also calculated separately within the WM and CSF masks. For each CompCor decomposition, the $k$

components with the largest singular values are retained, such that the retained components' time series are sufficient to explain 50 percent of variance across the nuisance mask (CSF, WM, combined, or temporal). The remaining components are dropped from consideration.

The head-motion estimates calculated in the correction step were also placed within the corresponding confounds file. The confound time series derived from head motion estimates and global signals were expanded with the inclusion of temporal derivatives and quadratic terms for each (Satterthwaite et al., 2013). Frames that exceeded a threshold of 0.5 mm FD or 1.5 standardized DVARS were annotated as motion outliers. Additional nuisance timeseries are calculated by means of principal components analysis of the signal found within a thin band (*crown*) of voxels around the edge of the brain, as proposed by Patriat et al. (2017).

The BOLD time-series were resampled into standard space, generating a *preprocessed BOLD run in MNI152NLin2009cAsym space*. All resamplings can be performed with *a single interpolation step* by composing all the pertinent transformations (i.e. head-motion transform matrices, susceptibility distortion correction when available, and co-registrations to anatomical and output spaces). Gridded (volumetric) resamplings were performed using antsApplyTransforms (ANTs), configured with Lanczos interpolation to minimize the smoothing effects of other kernels (Lanczos, 1964). Non-gridded (surface) resamplings were performed using mri_vol2surf (FreeSurfer).

Many internal operations of *fMRIPrep* use *Nilearn* 0.10.1 (Abraham et al., 2014; RRID:SCR_001362), mostly within the functional processing workflow. For more details of the pipeline, see the section corresponding to workflows in fMRIPrep's documentation (https://fmriprep.readthedocs.io/en/latest/workflows.html).

### Appendix B: Pre-specified Robustness Criteria

The sensitivity analyses reported in this study were governed by decision rules fixed before the corresponding models were estimated. These criteria were specified before those analyses were carried out, so that a preserved, weakened, or reversed effect could not be re-interpreted once its outcome was known. This appendix records the criteria; the outcomes of applying them are reported in the Results and Discussion.

#### Rationale

The originally specified first-level GLM time-locked the duration modulator to trial onset (the *trial-onset GLM*). Two model-specification concerns motivated a re-aligned GLM: (i) the duration modulator was displaced from the onset of the coherent target, and (ii) the trial-onset GLM contained no explicit response-related regressor. The *target-onset GLM* re-aligns the duration modulator to coherent-target onset and adds a response-prompt regressor (Methods, *First-level general linear models*), addressing both. Because any change to model specification unavoidably perturbs effect sizes, the decision rules below were fixed before the target-onset GLM was estimated. As set out in the Methods, the target-onset GLM is the better-specified model and is foregrounded in the revised manuscript; the criteria here governed how the two alignments were to be compared, and how the originally reported SMA interaction was to be interpreted in light of that comparison.

#### SMA context × duration interaction

The originally reported neural finding (trial-onset GLM) was a context × duration interaction in the supplementary motor area (SMA/pre-SMA): the $AF_{Dur} - DF_{Dur}$ contrast (AF parametric duration minus DF parametric duration), evaluated as the mean contrast estimates within the a priori SMA ROI (8-mm sphere at MNI 0, 12, 50; Wiener et al., 2010). Each rule below is applied to subject-level contrast estimates extracted within this ROI from the target-onset GLM.

| Outcome | Rule | Consequence for interpretation |
|---|---|---|
| **Preserved** | Sign of the SMA $AF_{Dur}$ − $DF_{Dur}$ contrast estimates matches the trial-onset GLM, and the ROI-level one-sample $z \geq 2.3$ (test against zero, two-tailed) | The SMA context × duration interaction is retained as a confirmed finding; the target-onset GLM is reported as a confirmatory sensitivity analysis. |
| **Weakened-but-directional** | Sign matches the trial-onset GLM, but the ROI-level $z < 2.3$ | The finding is reframed as context-modulated duration- or expectation-related activity in SMA, and the precise duration interpretation is treated as suggestive. |
| **Failed** | Sign reverses, or $\|z\| < 1$ (no directional signal) | The Results are reorganized around the behavioral PSE shift and descriptive ROI patterns; the duration-related interpretation is treated as exploratory. |

*Realized outcome:* under the target-onset GLM the SMA contrast estimates reversed sign, a *Failed* outcome. Accordingly, the revised manuscript does not advance a region-specific SMA duration claim; SMA/pre-SMA is reported as contributing to the network-level Context × Region effect, and the foregrounded results centre on the network-level pattern and the left-insula simple effect.

#### Context × region interaction (2 × 5 ANOVA)

In the trial-onset GLM the context × region interaction was strong ($F(4, 92) = 5.90$, $p_{GG} = .001$, Greenhouse–Geisser–corrected, $\eta^2_p = .20$). The rule below is applied to the same interaction re-estimated on target-onset-GLM contrast estimates.

| Outcome | Rule | Consequence for interpretation |
|---|---|---|
| **Preserved** | Interaction $p_{GG} < .01$ and $\eta^2_p \geq .10$ | The context-dependent regional pattern is retained as reported. |

| Outcome | Rule | Consequence for interpretation |
|---|---|---|
| **Weakened-but-directional** | $p_{GG}$ between .01 and .05, or $\eta^2_p$ between .06 and .10 | The claim is softened to a context-dependent regional weighting. |
| **Failed** | $p_{GG} > .05$, or $\eta^2_p < .06$ | The regional-dissociation claim is dropped, and only the ROI-level simple effects are reported, descriptively. |

#### Holm-corrected AF-versus-DF simple effects

The five bilateral-averaged a priori ROIs (SMA, putamen, insula, cerebellum VI, IFG) are each tested for an AF-versus-DF difference with a paired *t*-test, Holm-corrected across the five tests. A region is taken to show a reliable simple effect only if its Holm-corrected $p < .05$. The directional expectations fixed in advance were AF > DF for SMA, putamen, and IFG; DF > AF for the insula; and no directional prediction for cerebellum VI.

#### gPPI connectivity

Because the parametric and non-parametric PPI interaction terms differ intrinsically in detectability, the gPPI analysis is reported as exploratory and convergent, never as confirmatory.

| Pattern | Rule | Consequence for interpretation |
|---|---|---|
| **Convergent** | A parametric $AF_{Dur} - DF_{Dur}$ PPI cluster at $z > 3.1$, $k > 10$ in the SMA-seed maps, with parametric-versus-non-parametric PPI-term correlations $\|r\| < .5$ | The gPPI result is retained in the main text, with per-subject diagnostics in Appendix E. |
| **Direction-dependent** | A cluster in only one contrast direction | Both directions are reported explicitly with *z* and *k*, and the asymmetry is flagged in the Discussion. |

| Pattern | Rule | Consequence for interpretation |
|---|---|---|
| **Regressor-structure artifact** | PPI-term correlations $\|r\| \geq .5$, or the effect disappears under the a priori SMA-ROI seed | The full gPPI is moved to Appendix E, and the main text retains only a one-sentence exploratory mention. |

#### Weber-fraction equivalence

To address whether temporal precision differs between contexts, the AF-versus-DF Weber-fraction difference is evaluated against the rules below.

| Outcome | Rule | Consequence for interpretation |
|---|---|---|
| **Equivalent** | TOST $p < .05$ against equivalence bounds of ±0.5 × the auditory-bisection effect size of Zhu et al. (2021) | Precision is reported as statistically equivalent across contexts — a criterion shift rather than a precision change. |
| **Indeterminate** | TOST $p \geq .05$ and the frequentist test $p \geq .05$ | No precision difference is detected, but the evidence is reported as indeterminate. |
| **Underpowered** | Bayes factor $BF_{01} < 3$ | The Bayes factor is reported, with explicit acknowledgement of limited evidence either way. |

#### Behavioral psychometric refit

Psychometric functions are refitted with a symmetric lapse parameter. The $\lambda \leq .10$ fit serves as the headline model and the $\lambda \leq .05$ fit as a stricter sensitivity check; the headline bound was set so that few per-condition fits sat at the lapse boundary.

### Appendix C: Behavioral Supplementary Analyses

#### Psychometric Lapse-Model Sensitivity

The behavioral analysis fits a cumulative-Gaussian psychometric function with a symmetric lapse parameter $\lambda$ bounded at .10. To confirm that the context effect on the point of subjective equality (PSE) is not an artifact of this modeling choice, we refit the psychometric function under four lapse specifications: the original lapse-free model ($\lambda$ fixed at 0), two bounded models ($\lambda \leq .05$ and $\lambda \leq .10$), and an unbounded model in which $\lambda$ was freely estimated over its full admissible range, [0, 0.5]. All models were fit by binomial maximum likelihood separately for each participant and context ($N = 24$).

The AF − DF PSE shift is stable across all four specifications, ranging only from 148 to 152 ms (all $t(23) > 6.9$, $p < .001$; Table C1), and no specification yields a reliable AF − DF Weber-fraction difference. In the unbounded fit the freely estimated lapse rate stayed low (mean $\lambda = .043$) and no condition fit reached the 0.5 boundary, confirming that the bounded models were not unduly constrained. The 19 of 48 condition fits that sat at the boundary under $\lambda \leq .05$ motivated the more permissive $\lambda \leq .10$ headline bound, at which only 8 of 48 fits remained at the boundary.

**Table C1. Context effect on the PSE across four psychometric lapse-model specifications ($N$ = 24).** The PSE shift is AF − DF with its 95% confidence interval; Weber values are context means; $t$ and $p$ are paired tests across participants.

| Lapse model | Mean $\lambda$ | PSE shift (ms) [95% CI] | $t(23)$ | $p$ | Weber AF / DF | Weber $p$ |
|---|---|---|---|---|---|---|
| Lapse-free ($\lambda = 0$) | 0 (fixed) | 148 [104, 192] | 6.95 | < .001 | .216 / .201 | .412 |
| Bounded, $\lambda \leq .05$ | .026 | 152 [107, 197] | 7.04 | < .001 | .197 / .180 | .359 |
| Bounded, $\lambda \leq .10$ (headline) | .037 | 151 [106, 195] | 6.93 | < .001 | .179 / .166 | .476 |
| Unbounded ($\lambda$ free in [0, .5]) | .043 | 150 [105, 194] | 6.96 | < .001 | .172 / .164 | .672 |

### Appendix D: Supplementary fMRI Analyses

#### D.1 Design-Matrix Collinearity Diagnostics

For the primary (trial-onset) and target-onset GLMs we computed, per participant and run, the pairwise Pearson correlations among the task regressors and the variance inflation factor (VIF) of each task regressor after regressing it on all other non-constant design columns (48 runs per GLM). Table D1 summarizes the maxima across all runs.

The parametric duration modulators, the regressors carrying the effects of interest, are well-conditioned in both GLMs: their VIFs do not exceed 1.16 and their correlations with the corresponding context boxcar stay below $|r| = 0.26$. The target-onset GLM shows a higher overall maximum VIF (2.17) and maximum correlation ($|r| = 0.70$), but this is confined to the structural correlation between the context boxcar and the response-prompt stick regressor, which sits at a fixed 2.5-s offset from the boxcar; it does not involve the duration modulators. All VIFs remain far below the conventional VIF > 5 threshold for collinearity concern.

**Table D1. Design-matrix collinearity diagnostics: maxima across 48 runs per GLM.**

| GLM | Max task-regressor VIF | Max $|r|$, task regressors | VIF, AF / DF duration modulator | $|r|$, modulator vs. its boxcar (AF / DF) |
|---|---|---|---|---|
| Primary (trial-onset) | 1.22 | 0.21 | 1.13 / 1.12 | 0.21 / 0.18 |
| Target-onset | 2.17 | 0.70* | 1.16 / 1.16 | 0.23 / 0.26 |

*The target-onset maximum correlation is the context-boxcar–response-prompt structural correlation; modulator-related correlations remain ≤ 0.26.

#### D.2 Lateralized ROI Follow-Up

The main-text 2 × 5 Context × Region ANOVA and the Holm-corrected simple-effect tests (Table 2) average each bilateral ROI across hemispheres. As a descriptive check on that averaging, Table

D2 reports AF-versus-DF paired *t*-tests at each ROI separately by hemisphere, on primary-GLM contrast estimates (uncorrected for multiple comparisons).

The lateralized pattern mirrors the bilateral-averaged results and does not reveal a hemisphere-specific effect that averaging obscures. The two insula ROIs show the same DF > AF direction (right $p = .027$, left $p = .058$), and SMA/pre-SMA, a single midline ROI, shows the AF > DF effect already reported in Table 2. Putamen, cerebellum VI, and IFG show no consistent lateralized effect. Hemispheric averaging is therefore descriptively justified.

**Table D2. Lateralized AF-versus-DF paired tests at each a priori ROI (primary GLM, *N* = 24, uncorrected).** Contrast estimates are group means.

| ROI (hemisphere) | AF | DF | AF − DF | *t*(23) | *p* | *d* |
|---|---|---|---|---|---|---|
| SMA/pre-SMA (midline) | 13.25 | 1.18 | 12.07 | 2.61 | .016 | 0.53 |
| Putamen (L) | 1.16 | −0.92 | 2.08 | 0.63 | .533 | 0.13 |
| Putamen (R) | 2.97 | −0.49 | 3.46 | 0.96 | .347 | 0.20 |
| Insula (L) | −3.02 | 5.56 | −8.58 | −1.99 | .058 | −0.41 |
| Insula (R) | −3.09 | 5.47 | −8.56 | −2.36 | .027 | −0.48 |
| Cerebellum VI (L) | 3.14 | −0.40 | 3.53 | 1.14 | .267 | 0.23 |
| Cerebellum VI (R) | 0.97 | 0.70 | 0.28 | 0.08 | .935 | 0.02 |
| IFG (L) | −0.27 | 6.67 | −6.94 | −1.65 | .112 | −0.34 |
| IFG (R) | 9.24 | 3.44 | 5.81 | 1.89 | .071 | 0.39 |

#### D.3 Trial-Onset GLM SVC (Comparison)

Table D3 reports small-volume-corrected peak statistics for the trial-onset GLM — the originally registered analysis, using the identical SVC method applied to the foregrounded target-onset GLM in Table 1. Under trial-onset alignment, duration-related modulation loaded on the AF (long-biased) context, surviving SVC in SMA/pre-SMA, right putamen, and right IFG, and the Context × Duration interaction peaked in SMA/pre-SMA in the AF > DF direction. Under target-onset alignment (Table 1) the same analyses instead localize to the DF (short-biased) context and to the right insula. Because target duration and target-onset latency are complementary cues confounded by the fixed display window, the regional attribution of duration-related signal — and the sign of the SMA interaction — depends on where the duration modulator is placed; this alignment-sensitivity is the limitation discussed in the main text.

**Table D3. Trial-onset GLM: SVC peak z-statistics (FWE-corrected p-values) within the 8-mm spherical a priori ROIs. Bold indicates *p*FWE-SVC < .05; conventions as in Table 1.**

| ROI | TimeDur *z* (*p*) | AF_TimeDur *z* (*p*) | DF_TimeDur *z* (*p*) | Context × Duration *z* (*p*) |
|---|---|---|---|---|
| SMA | 3.26 (.083) | **4.52 (<.001)** | 1.36 (1.00) | **3.45 (.041)** |
| R Putamen | 2.85 (.302) | **3.41 (.045)** | −1.35 (1.00) | 2.54 (.763) |
| L Putamen | 1.51 (1.00) | 1.57 (1.00) | −1.67 (1.00) | 2.75 (.401) |
| R IFG | 3.16 (.116) | **3.79 (.011)** | 2.64 (.613) | 2.88 (.290) |
| L IFG | 2.69 (.520) | −1.50 (1.00) | 3.25 (.084) | −2.48 (.951) |
| R Insula | 1.92 (1.00) | −2.99 (.200) | 2.46 (.993) | −2.96 (.223) |
| L Insula | 1.64 (1.00) | −2.68 (.538) | **3.42 (.046)** | −3.26 (.082) |
| R Cerebellum VI | **3.45 (.042)** | 2.72 (.486) | 2.57 (.754) | −1.48 (1.00) |
| L Cerebellum VI | 1.89 (1.00) | 2.99 (.215) | −1.56 (1.00) | 2.58 (.753) |

### Appendix E: gPPI Supplementary Results

The generalized psychophysiological interaction (gPPI) analysis is reported as exploratory (Methods, *Generalized Psychophysiological Interaction Analysis*). It was run from two seeds: the data-driven AFTimeDur − DFTimeDur whole-brain peak (MNI 6, −4, 58) and the a priori SMA/pre-SMA ROI (MNI 0, 12, 50), each an 8-mm sphere, and computed for both the parametric interaction contrast ($PPI_{AFTimeDur} - PPI_{DFTimeDur}$) and the non-parametric interaction contrast ($PPI_{AF} - PPI_{DF}$), in both directions. All 24 participants were processed successfully for both seeds.

#### E.1 Whole-Brain Cluster Peaks

The non-parametric context PPI contrast produced no cluster surviving $z > 3.1$, $k > 10$ in either direction or either seed. The parametric PPI contrast produced clusters in both directions and both seeds (AF > DF: 10 clusters from the data-driven seed, 17 from the a priori seed; DF > AF: 7 and 6 clusters, respectively). The strongest peaks converged across seeds on the same partners (Table E1): right IFG/pre-motor cortex for AF > DF and right V5/MT+ for DF > AF.

**Table E1. Strongest gPPI peaks for the parametric interaction contrast, both seeds and directions ($z > 3.1$, $k > 10$).**

| Contrast | Seed | Peak MNI (x, y, z) | *z* |
|---|---|---|---|
| AF > DF | Data-driven (6, −4, 58) | 30, 12, 50 | 4.85 |
| | | −6, −54, 11 | 4.45 |
| | | −42, −82, 34 | 4.29 |
| | A priori SMA (0, 12, 50) | 30, 12, 50 | 4.67 |
| | | −42, −82, 34 | 4.60 |
| | | 50, 36, 30 | 4.40 |
| DF > AF | Data-driven (6, −4, 58) | 42, −70, −6 | 5.22 |
| | | −42, −76, 1 | 4.99 |
| | | −30, −90, 7 | 4.85 |
| | A priori SMA (0, 12, 50) | 42, −70, −6 | 5.00 |
| | | −42, −76, 1 | 4.96 |
| | | −24, −84, −13 | 4.59 |

#### E.2 ROI-Level Coupling

We also extracted the mean PPI contrast estimate within each of the five a priori timing ROIs. Table E2 reports the AF-versus-DF coupling difference for the parametric contrast at each seed (paired $t$-tests, $N = 24$, uncorrected). No ROI-to-ROI coupling difference survived FDR correction; the strongest effects were a sub-threshold SMA and insula coupling reduction under the data-driven seed.

**Table E2. ROI-level gPPI coupling, parametric AF − DF contrast ($N = 24$, uncorrected).**

| ROI | Data-driven seed *t* (*p*) | A priori SMA seed *t* (*p*) |
|---|---|---|
| SMA | −1.82 (.083) | −1.39 (.179) |

| ROI | Data-driven seed *t* (*p*) | A priori SMA seed *t* (*p*) |
|---|---|---|
| Putamen | −1.46 (.157) | −1.08 (.293) |
| Insula | −1.72 (.099) | −1.20 (.242) |
| IFG | −0.55 (.589) | 0.18 (.861) |
| Cerebellum VI | −0.27 (.787) | 0.61 (.551) |

#### E.3 Per-Subject PPI-Term Diagnostics

Because the parametric and non-parametric PPI interaction terms differ intrinsically in detectability, we computed per-subject diagnostics of both terms, the L2 norm, the temporal variance, and the within-context correlation between the parametric and non-parametric PPI regressors, to confirm that any detectability asymmetry reflects regressor structure rather than neural connectivity. Across the 24 participants the parametric–non-parametric correlation was modest in both seeds (maximum $|r| = 0.32$, mean = 0.30), well below the 0.5 group-level threshold specified in Appendix B. No participant was flagged under the data-driven seed; one participant (sub-020) was flagged under the a priori seed for a high-variance PPI term, with clean correlation values, and was retained. No participant was excluded.
